# Integrative Clinical–Molecular Modeling Identifies *LRRN4CL* as a Determinant of Structural and Functional Myocardial Improvement

**DOI:** 10.64898/2026.04.21.720029

**Authors:** Ezra Johnson, Joseph R. Visker, Ben J. Brintz, Christos P. Kyriakopoulos, James Jeong, Yuyu Zhang, Thirupura S. Shankar, Yanni Hillas, Iosif Taleb, Rachit Badolia, Junedh M. Amrute, Chris J. Stubben, Luis Cedeno-Rosario, Ioannis Kyriakoulis, Konstantinos Sideris, Jing Ling, Rana Hamouche, Eleni Tseliou, Sutip Navankasattusas, Gregory S. Ducker, Jared Rutter, William L. Holland, Scott A. Summers, TingTing Hong, Steven C. Koenig, Thomas C. Hanff, Kory J. Lavine, Tom Greene, Stephen Bailey, Rami Alharethi, Craig H. Selzman, Palak Shah, Hongchao Guo, Mark S. Slaughter, Manreet K. Kanwar, Stavros G. Drakos

## Abstract

**Background:** Mechanical ventricular unloading and systemic circulatory support with left ventricular assist devices (LVADs) enable myocardial recovery in a subset of advanced heart failure (HF) patients, but predictors and mechanisms of recovery are not well understood. Integrating clinical and molecular data may improve identification of patients most likely to recover and uncover biologically relevant targets in HF.

**Methods:** We collected and analyzed left ventricular apical myocardial tissue and clinical data from 208 patients undergoing LVAD implantation across five centers. Pre-implant transcriptomic profiles (22,373 mRNA transcripts) were integrated with 59 clinical variables using supervised machine learning with repeated cross-validation to identify and prioritize features associated with myocardial recovery, defined as a binary outcome based on improvement in left ventricular ejection fraction (LVEF ≥40%) and left ventricular end-diastolic diameter (LVEDD ≤5.9 cm). We also modeled functional (LVEF) and structural (LVEDD) improvement as a continuous outcome without any predefined LVEF and LVEDD pathological thresholds. Feature prioritization was followed by validation in human myocardial tissue and mechanistic interrogation in human induced pluripotent stem cell-derived cardiomyocytes (iPSC-CMs).

**Results:** Integrative models achieved modest discrimination for myocardial recovery as a binary categorical outcome (maximum mean cross-validated area under the curve 0.73±0.15), identifying clinical features such as HF duration, LVEDD, HF pharmacologic therapy, and device configuration. Leucine-rich repeat neuronal 4C-like (*LRRN4CL*), measured in human myocardium, consistently emerged as a top transcriptomic predictor across both binary and continuous metric models (functional and structural). Higher pre-LVAD *LRRN4CL* expression was associated with reduced likelihood of myocardial recovery and localized primarily to cardiomyocytes. In iPSC-CMs, *LRRN4CL* overexpression localized to the sarcoplasmic reticulum, induced transcriptional remodeling characterized by suppression of contractile pathways and activation of stress programs, impaired calcium handling, impaired contraction–relaxation kinetics, and diminished mitochondrial respiratory reserve capacity.

**Conclusions:** Integration of clinical and myocardial transcriptomic data identifies *LRRN4CL* as a novel marker associated with impaired myocardial recovery following LVAD-mediated ventricular unloading and systemic circulatory support. These findings move beyond predictive modeling, linking integrative computational discovery to cardiomyocyte dysfunction and providing a translational framework for biologically informed risk stratification and therapeutic targeting for myocardial recovery.

**CLINICAL PERSPECTIVE:** *What Is New?:*

- Integrative clinical and myocardial transcriptomic modeling identifies *LRRN4CL* as a novel molecular determinant of structural and functional changes after LVAD-mediated ventricular unloading and enhanced systemic circulatory support.
- Elevated *LRRN4CL* expression is associated with adverse remodeling signatures, impaired calcium handling, and stress responses in human iPSC-derived cardiomyocytes.
- Experimental overexpression of *LRRN4CL* directly disrupts calcium cycling, contractile performance, and mitochondrial respiration linking molecular signature to functional phenotype.

*What Are the Clinical Implications?:*

- Identification of *LRRN4CL* as a marker associated with impaired myocardial recovery supports future efforts toward biologically informed risk stratification for patients undergoing LVAD therapy.
- *LRRN4CL* as a marker of cardiac improvement potential may extend beyond advanced HF to earlier stage disease patients and inform prognosis, risk stratification, and response to medical therapies.
- These findings highlight *LRRN4CL*-associated pathways as potential therapeutic targets and demonstrate how integrative clinical–transcriptomic approaches can move beyond clinical prediction toward identification of new biologically precise therapeutic targets in HF following a bedside to bench and back approach.

## INTRODUCTION

Heart failure (HF) is a progressive, chronic disease associated with high morbidity and mortality (1). Left ventricular assist devices (LVADs) are an established therapeutic approach for advanced HF patients with refractory symptoms despite guideline-directed medical therapy (2). LVAD therapy offers a unique investigational model by providing left ventricular (LV) apical myocardial tissue at the time of implantation. LVADs, through enhanced systemic circulatory perfusion and volume and pressure unloading of the failing LV, can enable structural and functional cardiac improvement (3–9). On the molecular level, LVADs result in positive adaptations to cardiomyocyte metabolism, excitation–contraction coupling, and gene expression (10–15). Although certain clinical characteristics (6,8,9,16) and biological observations (12,14,17,18) have been associated with a higher tendency for structural and functional cardiac improvement, the interplay between clinical outcomes and underlying biological phenomena driving these changes is not well understood. Currently, there is a lack of accurate predictive markers and molecular mechanisms to forecast LVAD-mediated myocardial recovery among advanced HF patients and innovative bioinformatic approaches are needed (19,20).

Analyses that combine clinical and molecular profiles using machine learning (ML) have proven useful for the diagnosis, prognosis, and classification of other human diseases (21,22). Compared to traditional regression-based models, ML can integrate high-dimensional clinical and molecular data to uncover biologically informative patterns that may not be apparent using traditional pre-specified models, particularly for complex diseases such as HF.

Recent studies have leveraged molecular profiling to elucidate mechanisms of myocardial recovery (23,24,25); however, direct coupling of predictive modeling with experimental validation remains limited. We hypothesized that integrative analysis of pre-LVAD clinical characteristics and transcriptomic profiles would identify predictors of LVAD-mediated myocardial recovery. Specifically, we posited that combining molecular and clinical data using supervised ML would identify biologically meaningful features associated with both functional and structural cardiac improvement. To evaluate this, we coupled ML-based feature prioritization with experimental validation in human iPSC-CMs and cardiac tissue. Here, we link predictive modeling with mechanistic insight into myocardial adaptation in the field of LVAD-mediated myocardial recovery.

## METHODS

### Data Availability

Data are available from the corresponding author upon reasonable request in accordance with institutional human subject protections. First authors had full access to all study data and take responsibility for its integrity and analysis. Detailed methods are provided in the **Supplemental Materials**, including experimental procedures, analytic pipelines, and additional ML methodological details. Datasets have been deposited in the Gene Expression Omnibus (GEO) under accession numbers **GSE328059** and **GSE327946**. Code and associated resources are available at doi.org/10.5281/zenodo.19476279.

### Study Approval

The study was approved by institutional review boards at all participating centers, and all patients provided written informed consent. Human myocardial tissue collection was conducted under approved IRB protocols (IRB_00030622 and IRB_00061825).

### Study Population

Patients with advanced HF undergoing continuous-flow LVAD implantation across five U.S. centers were included (see details in Supplemental Material). Patients with acute etiologies of HF, hypertrophic or infiltrative cardiomyopathies, baseline LVEF ≥40%, inadequate follow-up (<3 months), or early transplantation/death were excluded.

### Clinical Data Collection and Echocardiography

Demographics, clinical characteristics, laboratory data, hemodynamics, and imaging data were collected. Echocardiography was performed at baseline and serially after LVAD implantation per institutional protocols and guideline-recommended standards. Patients achieving an LV ejection fraction (LVEF) ≥40% and LV end-diastolic diameter (LVEDD) ≤5.9 cm at any time within one year following LVAD implantation were classified as *responders (R);* all others were designated *non-responders (NR).* In addition to **binary** classification, changes in LVEF and LVEDD were analyzed as **continuous metrics** of functional and structural recovery without using any predefined thresholds (**Figure 1B**).

**Figure 1.**
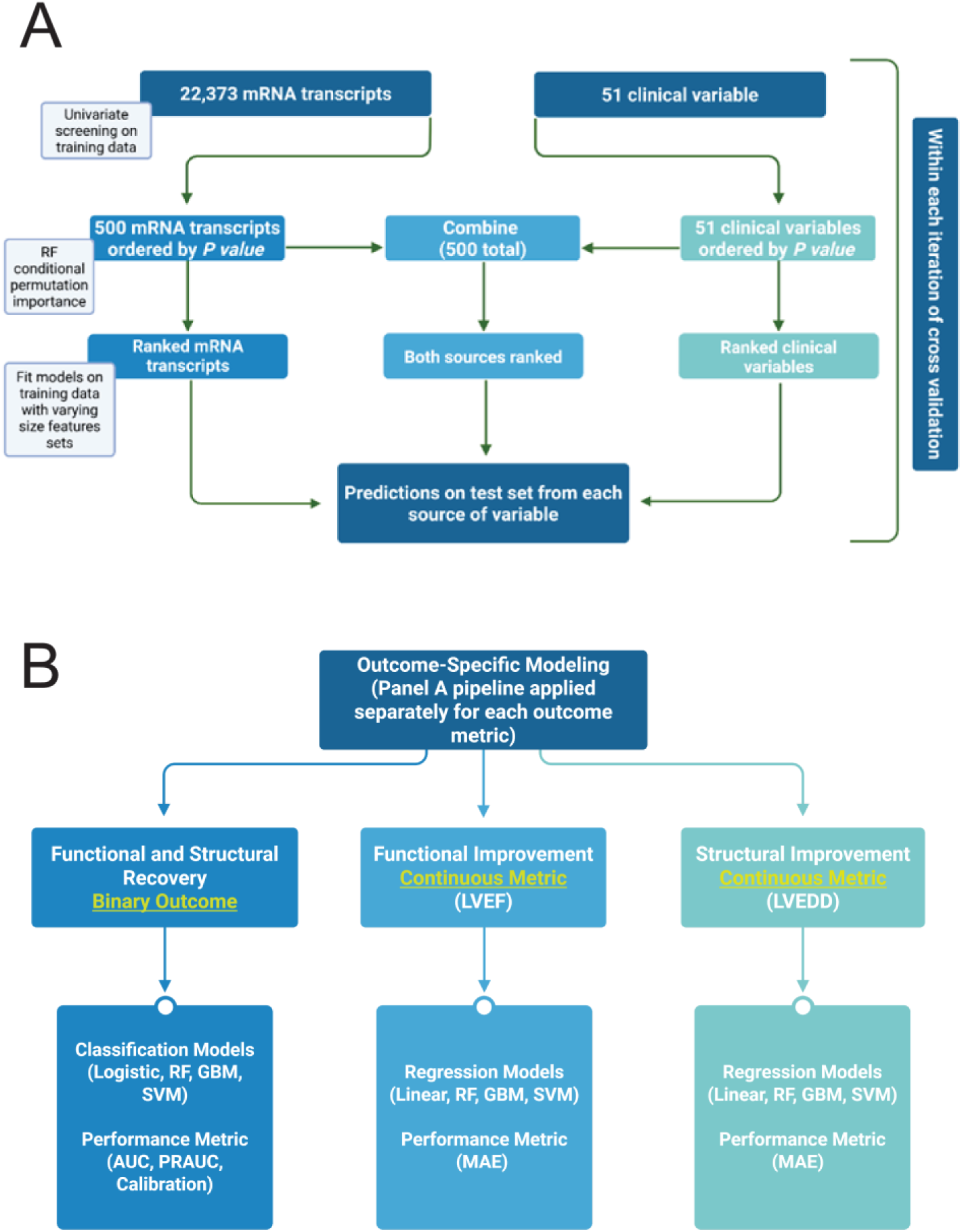
Machine learning pipeline for feature screening, selection, and outcome modeling. (A) mRNA transcripts (n=22,373) and clinical variables (n=51) were independently screened within cross-validation using univariate analysis, with top features advanced to machine learning–based ranking using random forest importance measures. Feature subsets of varying sizes were evaluated within cross-validation, and combined feature sets were used to generate predictions on held-out test sets. (B) The pipeline shown in panel A was applied independently for each outcome metric within cross-validation, incorporating outcome-specific univariate screening and feature ranking. Three complementary outcomes were modeled: **binary myocardial recovery** (responder vs non-responder), functional recovery assessed by **continuous post-LVAD LVEF**, and structural recovery assessed by **continuous post-LVAD LVEDD**. Outcome-specific modeling approaches and performance evaluation metrics are shown.

### Human Myocardial Tissue Procurement

Left ventricular apical core tissue was collected at the time of LVAD implantation and immediately flash-frozen as previously described (14). Non-failing donor myocardial tissue from hearts unsuitable for transplantation due to non-cardiac reasons (infection, donor-recipient size mismatch, etc.) served as controls. Molecular analyses were performed on pre-LVAD samples only.

### RNA Sequencing and Transcriptomic Analysis

Bulk RNA sequencing was performed on LV myocardial tissue and iPSC-derived cardiomyocytes. Reads were aligned to the human reference genome (GRCh38), and differential expression analysis was conducted using DESeq2 with batch correction. For human data, transcriptomic validation was performed using RT-qPCR. For iPSC-CMs, pathway enrichment analyses were conducted using FGSEA against MSigDB pathway databases. Detailed library preparation, sequencing, alignment parameters, filtering criteria, and enrichment methods for human tissue and iPSC-CMs are provided in the **Supplemental Materials**.

### Predictive Modeling and Machine Learning–Based Feature Prioritization

Predictive analyses integrated pre-LVAD transcriptomic (22,373 transcripts) and clinical (59 variables) features. Clinical variables with ≥40% missingness were excluded; remaining missing data were handled using multiple imputation with chained equations. Given the high dimensionality of transcriptomic data, feature selection employed a two-stage approach consisting of initial univariate regression-based screening followed by machine learning–based variable importance ranking to prioritize candidate predictors. Feature prioritization and model performance were evaluated using repeated 90/10 cross-validation (100 iterations). Models assessed included logistic regression, random forest, gradient boosting, and support vector machines. Model performance was evaluated using AUC and precision-recall curves (PRAUC) for binary outcomes and mean absolute error (MAE) for continuous outcomes (26). Models were designed for biological feature discovery rather than clinical deployment.

### iPSC-CM Differentiation and Viral Manipulation

Human iPSCs were differentiated into cardiomyocytes using a chemically defined Wnt-modulation protocol (27,28). Cardiomyocytes were metabolically purified prior to experimentation. *LRRN4CL* overexpression was achieved using AAV9 or adenoviral vectors. Control viruses were used in parallel.

### Contractility Assays using iPSC-CMs

Spontaneous contraction recordings were obtained by brightfield video microscopy. *LRRN4CL* overexpression and GFP control groups were analyzed using CONTRACTIONWAVE motion-tracking software (29,30) to quantify contraction-relaxation kinetics.

### Calcium Handling and Live-Cell Imaging in iPSC-CMs

Live-cell calcium imaging was performed using fluorescent calcium indicators (Cal-520 and X-Rhod1) following caffeine-stimulation (20mM) to assess calcium transient dynamics and sarcoplasmic reticulum function.

### Mitochondrial Bioenergetic Assessment by Seahorse Assay in iPSC-CMs

Oxygen consumption rate (OCR) was measured using the Seahorse XF Mito Stress Test. Parameters of mitochondrial respiration were calculated according to manufacturer guidelines and normalized to cell number.

### RNAscope and Immunofluorescence

Multiplex fluorescent RNAscope in situ hybridization was performed on formalin-fixed paraffin-embedded myocardial tissue and cultured iPSC-CMs, followed by immunofluorescence staining for cell-type markers.

### Statistical Analysis

Statistical analyses were conducted using R (v4.1.0) and GraphPad Prism (v9.01). Data are presented as mean±SD unless specified otherwise, and statistical significance was defined *a priori* as p<0.05. Statistical tests for individual experiments are described in the corresponding figure legends and detailed information is provided in the **Supplemental Materials**.

## RESULTS

### Patient Characteristics

Our study cohort comprised 208 patients with chronic advanced HF undergoing durable, continuous-flow LVAD implantation at the following participating institutions: the Utah Cardiac Recovery (UCAR) Program (n=123), Allegheny General Hospital (n=53), and the University of Louisville (n=32) (**Supplemental Table S1**). Baseline clinical characteristics stratified by recovery status are shown in **Table 1**. Compared with non-responders, responders were younger (51.4±18.3 vs. 57.1±13.7 years; p=0.042) and had a shorter duration of HF symptoms before LVAD implantation (43.5±60.2 vs. 80.4±72.9 months; p<0.001). Responders were also less likely to have pre-implant cardiac resynchronization therapy and/or an implantable cardioverter-defibrillator (73.1% vs. 87.0%; p=0.019, likely a surrogate of the shorter duration of HF symptoms) and had smaller baseline LV chamber size, as reflected by lower pre-implant LVEDD (6.6±1.0 vs. 7.0±1.2 cm; p=0.048). Baseline LVEF was not significantly different between responders and non-responders (19.0±7.6% vs. 17.1±6.5%, respectively; p=0.082). Most other baseline demographic characteristics, comorbidities, INTERMACS profile, pre-operative supportive therapies, hemodynamic measures, and laboratory values were generally similar between groups, except albumin which was modestly lower in responders (3.6±0.6 vs. 3.8±0.6 g/dL; p=0.020).

**Table 1:** Baseline clinical characteristics of LVAD responders and non-responders.

| Variable | Responders<br>(N=52) | Non-Responders<br>(N=156) | P-value |
| --- | --- | --- | --- |
| ***Demographics*** |  |  |  |
| Age, years | 51.4 (18.3) | 57.1 (13.7) | **0.042<sup>t</sup>** |
| Male sex, n (%) | 37 (71.2%) | 127 (81.4%) | 0.12<sup>c</sup> |
| Race |  |  | 0.39<sup>c</sup> |
| White, n (%) | 43 (82.7%) | 136 (87.2%) | - |
| Black, n (%) | 4 (7.7%) | 11 (7.1%) | - |
| Other, n (%) | 5 (9.6%) | 9 (5.8%) | - |
| Body Mass Index, kg/m<sup>2</sup> | 27.5 (5.1) | 29.2 (5.9) | 0.055<sup>t</sup> |
| Blood Type |  |  | 0.64<sup>f</sup> |
| A, n (%) | 13 (41.9%) | 33 (35.1%) |  |
| B, n (%) | 6 (19.4%) | 14 (14.9%) |  |
| AB, n (%) | 0 (0%) | 4 (4.3%) |  |
| O, n (%) | 12 (38.7%) | 43 (45.7%) |  |
| Rhesus D Antigen |  |  | 0.17<sup>c</sup> |
| Positive, n (%) | 39 (79.6%) | 120 (87.6%) |  |
| ***Medical History*** |  |  |  |
| Smoking, n (%) | 25 (48.1%) | 84 (54.5%) | 0.42<sup>c</sup> |
| Diabetes Mellitus, n (%) | 16 (30.8%) | 53 (34.4%) | 0.63<sup>c</sup> |
| Hypertension, n (%) | 26 (50%) | 86 (56.2%) | 0.44<sup>c</sup> |
| Ischemic Cardiomyopathy, n (%) | 15 (30%) | 56 (37.6%) | 0.33<sup>c</sup> |
| New York Heart Association Classification |  |  | **<0.001<sup>c</sup>** |
| 3, n (%) | 25 (48.1%) | 35 (22.6%) |  |
| 4, n (%) | 27 (51.9%) | 120 (77.4%) |  |
| Heart Failure Symptoms Duration, months | 43.5 (60.2) | 80.4 (72.9) | **<0.001<sup>t</sup>** |
| Cardiac Resynchronization/Implantable<br>Cardioverter Defibrillator, n (%) | 38 (73.1%) | 134 (87%) | **0.019<sup>c</sup>** |
| INTERMACS profile |  |  | 0.35<sup>f</sup> |
| 1, n (%) | 9 (23.1%) | 18 (13.4%) | - |
| 2, n (%) | 7 (17.9%) | 29 (21.6%) | - |
| 3, n (%) | 10 (25.6%) | 48 (35.8%) | - |
| 4-7, n (%) | 13 (33.4%) | 39 (29.1%) | - |
| Left Ventricular Assist Device Type |  |  | 0.31<sup>f</sup> |
| HeartMate II, n (%) | 34 (65.4%) | 89 (58.2%) | - |
| HeartMate 3, n (%) | 4 (7.7%) | 9 (5.9%) | - |
| HeartWare, n (%) | 10 (19.2%) | 45 (29.4%) | - |
| Other, n (%) | 4 (7.6%) | 10 (6.6%) | - |
| Left Ventricular Assist Device Configuration |  |  | 0.31<sup>c</sup> |
| Axial, n (%) | 36 (72%) | 97 (64.2%) |  |
| Left Ventricular Assist Device Support<br>Duration, days | 569.2 (623.5) | 564.0 (650.3) | 0.97<sup>t</sup> |
| ***Pre-operative Supportive Therapies*** |  |  |  |
| Inotrope Dependency, n (%) | 33 (66%) | 103 (68.7%) | 0.73<sup>c</sup> |
| Intra-aortic Balloon Pump, n (%) | 10 (20%) | 20 (13.2%) | 0.25<sup>c</sup> |
| Temporary Mechanical Circulatory Support, n (%) | 3 (6%) | 5 (3.3%) | 0.42<sup>f</sup> |
| **Heart Failure Medical Therapy** |  |  |  |
| Beta-Blockers, n (%) | 38 (74.5%) | 104 (68.4%) | 0.41<sup>c</sup> |
| Angiotensin Converting Enzyme Inhibitors/Angiotensin Receptor Blockers, n (%) | 33 (64.7%) | 86 (56.2%) | 0.29<sup>c</sup> |
| Mineralocorticoid Receptor Antagonists, n (%) | 20 (52.6%) | 86 (66.2%) | 0.13<sup>c</sup> |
| Diuretics, n (%) | 31 (81.6%) | 120 (93%) | 0.06<sup>f</sup> |
| **Right Heart Catheterization** |  |  |  |
| Right Atrial Pressure, mmHg | 11.5 (5.6) | 12.1 (6.6) | 0.53<sup>t</sup> |
| Pulmonary Capillary Wedge Pressure, mmHg | 23.3 (8.2) | 23.9 (8.4) | 0.62<sup>t</sup> |
| Cardiac Index, L/min/m<sup>2</sup> | 2.0 (0.7) | 1.9 (0.6) | 0.11<sup>t</sup> |
| **Laboratory Values** |  |  |  |
| White Blood Cells, x10<sup>9</sup>/μL | 9.1 (3.5) | 9.1 (3.7) | 0.98<sup>t</sup> |
| Platelets, x10<sup>9</sup>/μL | 218.9 (61.7) | 205.1 (72.6) | 0.23<sup>t</sup> |
| Hemoglobin, g/dL | 12.2 (2.3) | 12.6 (2.2) | 0.50<sup>t</sup> |
| Sodium, mEq/L | 135.2 (4.6) | 134.4 (4.9) | 0.36<sup>t</sup> |
| Blood Urea Nitrogen, mg/dL | 24.6 (15.7) | 27.8 (13.1) | 0.22<sup>t</sup> |
| Creatinine, mg/dL | 1.3 (0.6) | 1.3 (0.5) | 0.48<sup>t</sup> |
| Total Bilirubin, mg/dL | 1.5 (1.4) | 1.6 (1.1) | 0.58<sup>t</sup> |
| Albumin, g/dL | 3.6 (0.6) | 3.8 (0.6) | **0.020<sup>t</sup>** |
| B-type Natriuretic Peptide, pg/mL | 1633.4 (1721.6) | 1412.8 (2009.8) | 0.47<sup>t</sup> |
| **Echocardiographic Assessment** |  |  |  |
| Left Ventricular Ejection Fraction, % | 19.0 (7.6) | 17.1 (6.5) | 0.082<sup>t</sup> |
| Left Ventricular End-Diastolic Diameter, cm | 6.6 (1.0) | 7.0 (1.2) | **<0.048<sup>t</sup>** |

<sup>t</sup> T-test, <sup>c</sup> Chi-squared test, <sup>f</sup> Fisher's exact test.

### Machine Learning–Based Integration of Clinical and Transcriptomic Data

To identify pre-LVAD clinical and molecular predictors of myocardial recovery, we applied supervised machine learning integrating clinical variables and transcriptomic data. A total of 59 clinical variables and 22,373 mRNA transcripts were considered as candidate predictors. After exclusion of clinical variables with excessive missingness (≥40%) and imputation of remaining data, a two-stage feature selection framework was employed to prioritize informative predictors while limiting overfitting. This consisted of initial regression-based screening followed by random forest conditional permutation importance ranking.

Models were trained and evaluated using repeated cross-validation to assess predictive performance and feature stability. Three complementary outcome definitions were modeled: (**1**) **binary** myocardial recovery status (R or NR), (**2**) **functional improvement** assessed by post-LVAD LVEF as a continuous variable and metric, and (**3**) **structural improvement** assessed by post-LVAD LVEDD as a continuous variable and metric. This approach captures the spectrum of myocardial response to circulatory support and mechanical unloading and identifies transcriptomic features that predict recovery across phenotypes. **Figure 1** shows an outline of the ML pipeline, including the screening and testing methodology for the mRNA transcripts and clinical variables (**Figure 1A**), and the outcome modeling framework encompassing both binary and continuous metric (functional and structural) outcomes (**Figure 1B**). Comprehensive model performance metrics (**Supplemental Figures S5, S6, S8,** and **S9**), feature rankings (**Supplemental Table S2-4**, **Supplemental Figure S3**), calibration analyses (**Supplemental Figure S1**), and partial dependence plots (**Supplemental Figures S2, S7,** and **S10**) for each outcome are provided in the **Supplemental Methods** and further details are provided in the **Supplemental Results**.

### Machine Learning Prediction of LVAD-mediated Myocardial Recovery (binary outcome)

For prediction of LVAD-mediated myocardial recovery, defined as achieving both LVEF ≥40% and LVEDD ≤5.9 cm within one year of LVAD implantation, multiple supervised learning models were evaluated using clinical variables, transcriptomic variables, and their combination. Among integrative models, a gradient boosted machine (gbm) model incorporating 80 features achieved the highest discriminatory performance, with a mean cross-validated AUC of 0.731±0.15 (**Figure 2A**). Given the imbalance between R and NR, model performance was also assessed using PRAUC, which was maximized by a parsimonious 30-feature integrative model (PRAUC=0.445±0.253), outperforming the 80-variable model by 2% (PRAUC=0.425±0.242).

**Figure 2.**
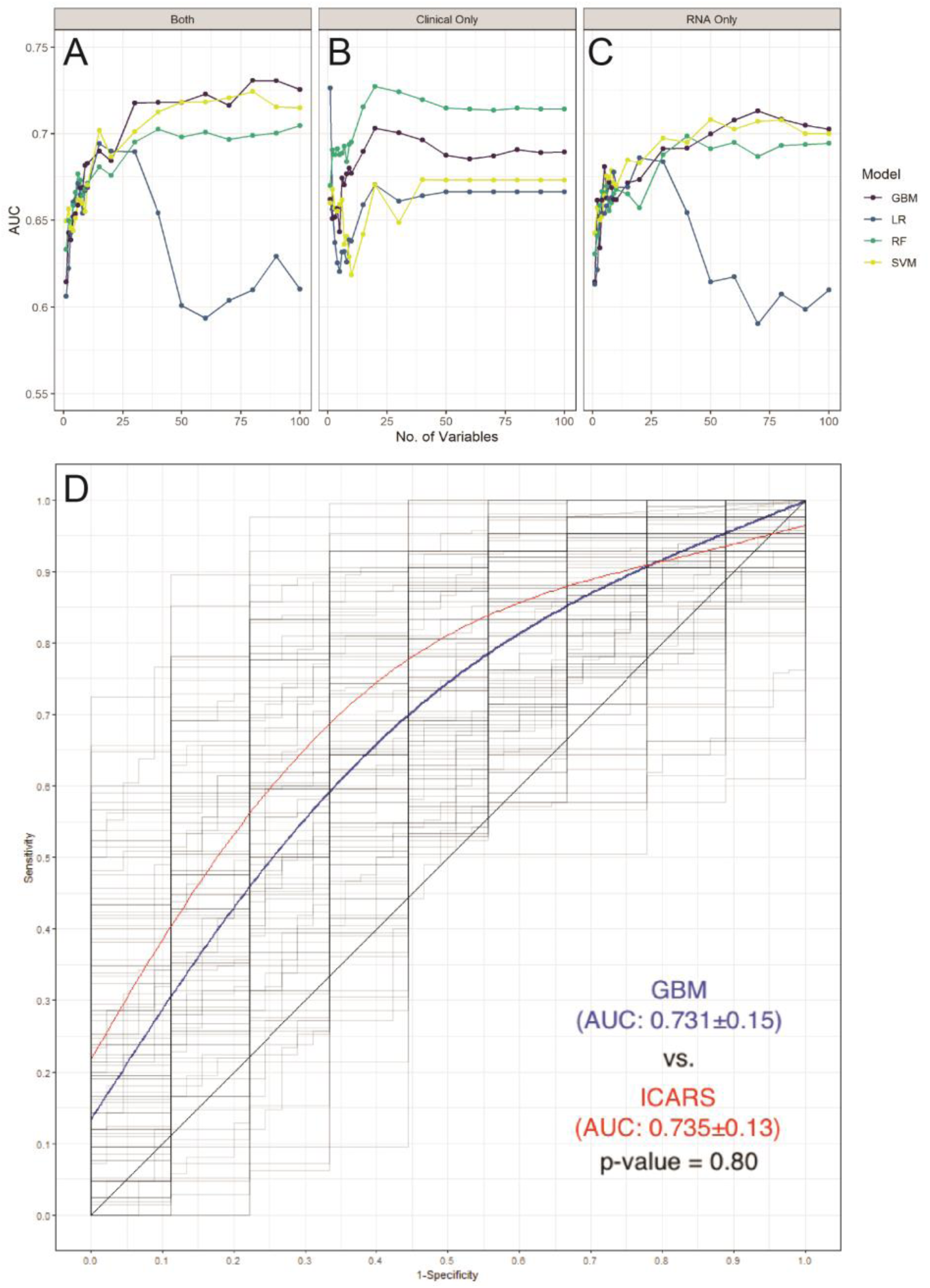
Cross-validated AUCs by (**A**) both clinical and mRNA variables, (**B**) only clinical variables, or **(C)** only mRNA variables and the number of variables used in the model for predicting binary LVAD-mediated myocardial recovery. Variables were ranked and selected using a two-stage screening process. (**D**) A smooth averaging of the ROC curves for the GBM model (solid blue line) with 80 variables, and the INTERMACS Cardiac Recovery (ICAR) score (solid red line) using cubic smoothing splines with 4 degrees of freedom as well as the ROC curve from each of the 100 iterations of cross-validation (shadowy lines). Gradient Boosted Regression (GBM: 5000 trees), Logistic Regression (LR), Random Forest (RF: 5000 trees), and Support Vector Machine (SVM: polynomial regression). A paired randomization test was used to compare GBM against ICARS resulting in a p-value of 0.80. ROC, Receiver Operator Characteristic.

Although the integrative gbm model achieved the highest mean AUC, clinical-only random forest (rf) models demonstrated more stable calibration across cross-validation iterations (**Supplemental Figure S1**), supporting their robustness. A random forest (rf) model with 20 clinical features achieved a mean cross-validated AUC of 0.727±0.14 and PRAUC of 0.471±0.237 (**Figure 2B**), while transcriptomic-only models achieved slightly lower discrimination (maximum AUC 0.713±0.14 and PRAUC of 0.388±0.227; **Figure 2C**). Comparing this to the previously published INTERMACS Cardiac Recovery (ICAR) score (9), which used a logistic regression model using 9 variables, yielded an AUC of 0.735±0.134, with no significant difference in performance relative to either integrative or clinical-only ML models (paired randomization; p-value=0.80 and p-value=0.49, respectively) (**Figure 2D**). This indicates that the ML approaches used here recapitulated established clinical predictors without exceeding their predictive ceiling.

Feature importance analysis from the binary recovery models identified established clinical predictors, including HF symptoms duration, baseline LV size/function, LVAD configuration, and HF pharmacologic therapy (**Supplemental Table S2**). Importantly, among transcriptomic predictors, leucine-rich repeat neuronal 4C-like (*LRRN4CL*) ranked among the top molecular features, with higher pre-LVAD expression associated with a lower probability of myocardial recovery. To further assess variable importance, we performed a univariate logistic regression analysis with each of the clinical and RNA variables as independent variables. HF symptoms duration (**Supplemental Figure S3A**), *ATF3*, and *LRRN4CL* (**Figure 3A**) appeared as significant (p<0.05) clinical and biological variables that were also top variables from the rf importance metrics.

**Figure 3.**
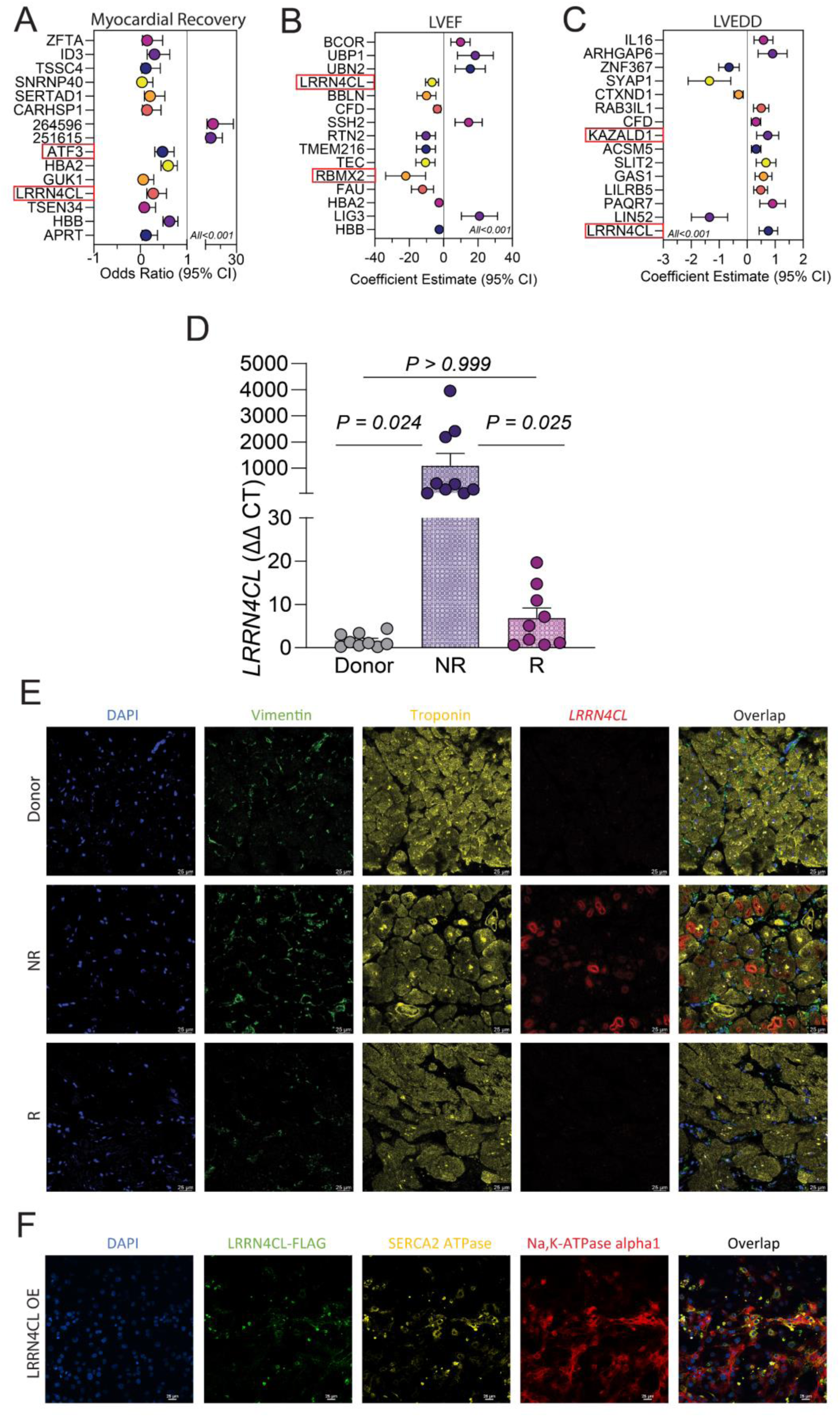
Identification and validation of *LRRN4CL* as a marker associated with myocardial recovery, cardiac function (LVEF), and structure (LVEDD). Forest plots show gene expression and clinical outcomes with coefficient estimates (95% CI) for the significantly identified mRNA transcripts associated with (**A**) improved binary myocardial recovery, (**B**) improved function (continuous LVEF), and (**C**) improved structure (continuous LVEDD), using linear regressions. Red boxes indicate variables that were identified via ML. (**D**) Relative gene expression of *LRRN4CL* in human myocardium as determined by Quantitative real-time PCR (RT-qPCR) (n=9). (**E**) Representative RNAscope and immunofluorescence images from non-failing donors, NR, and R samples showing DAPI (blue), vimentin protein (green), troponin protein (yellow), *LRRN4CL* RNA (red), and merged overlays, demonstrating cell-type localization and differential expression across conditions. Images are representative of three independent patient samples per group (n=3 per group: Donor, NR, and R), with three images obtained per sample. (**F**) Representative immunofluorescence images from human iPSC-CMs overexpressing LRRN4CL-FLAG (green), with DAPI (blue), SERCA2 ATPase (yellow), Na,K-ATPase alpha1 (red), and merged overlays. Images are representative of three independent samples (n=3), with three fields of view acquired per sample. Quantitative colocalization analysis of LRRN4CL-FLAG and SERCA2A revealed a mean Pearson’s correlation coefficient of 0.41±0.08, with Manders’ overlap coefficients of 0.557±0.067 (M1) and 0.592±0.071 (M2) (mean±SD, n=9), indicating moderate spatial colocalization between LRRN4CL-FLAG and SERCA2A. Costes randomization testing yielded P=1.0 in all replicates. Scale bars, 25 µm. LVEF indicates left ventricular ejection fraction; LVEDD, Left ventricular end-diastolic diameter; NR, non-responders; R, responders; OE, overexpression.

### Machine Learning Prediction of Functional Cardiac Improvement (continuous LVEF)

To model recovery as a *spectrum rather than a strictly binary outcome*, we modeled post-LVAD LVEF as a continuous variable. This approach enabled identification of predictors associated with degrees of functional improvement independent of categorical recovery thresholds.

Rf models using clinical variables alone achieved the best predictive performance for post-LVAD LVEF, with a 10-feature model yielding a mean cross-validated mean absolute error (MAE) of 9.93±1.71 (**Supplemental Figure S5**). The most influential clinical predictors included female sex, baseline LVEDD, LVAD type configuration, and baseline LVEF (**Supplemental Table S3**). These findings were consistent with known determinants of reverse functional remodeling during LVAD unloading and perfusion support. To validate, we performed a univariate linear regression analysis with each as an independent variable. LVEF, LVEDD, and HF symptoms duration (**Supplemental Figure S3B**) appeared as significant clinical variables across both linear regression and rf.

Among molecular predictors, *LRRN4CL* again emerged as a top-ranked feature. Higher pre-LVAD expression of *LRRN4CL* demonstrated a strong inverse association with post-LVAD LVEF improvement and ranked within the top 15 predictive transcripts (p<0.001; **Figure 3B**). Additional transcripts identified by the functional models included *RBMX2*, *EFCC1*, and *TXNL4B*; however, these transcripts did not demonstrate the same degree of consistency across recovery outcomes as *LRRN4CL*. Detailed rankings and analyses for other top ML-identified transcripts, including *RBMX2*, *EFCC1*, and *TXNL4B*, are provided in the **Supplemental Results** and **Supplemental Figure S4**.

### Machine Learning Prediction of Structural Cardiac Improvement (continuous LVEDD)

Structural myocardial improvement was evaluated by modeling post-LVAD LVEDD as a continuous variable. In these analyses, rf models using clinical variables again demonstrated the strongest performance, with a 6-feature model achieving a mean cross-validated MAE of 0.79±0.13 (**Supplemental Figure S8**). Key clinical predictors included female sex, baseline LVEDD, LVAD type, platelet count, and B-type natriuretic peptide levels (**Supplemental Table S4**).

Transcriptomic models identified several molecular predictors of structural reverse remodeling, including *TCIM* and *KAZALD1*. Notably, *LRRN4CL* again ranked among the top predictive transcripts (**Figure 3C, Supplemental Table S4**), with lower pre-LVAD expression significantly associated with greater reduction in LVEDD following LVAD support (p<0.001; **Figure 3C**).

### Convergence of machine learning models on *LRRN4CL* motivates functional validation

Across all models, *LRRN4CL* emerged as a consistent transcriptomic predictor of myocardial recovery from LVAD unloading and circulatory support. This finding across multiple analyses distinguished *LRRN4CL* from other potential transcripts and justified further experimental validation.

### *LRRN4CL* Is Upregulated in Non-Responders and Localizes to Cardiomyocytes in Failing Human Myocardium

To validate the association between *LRRN4CL* expression and myocardial recovery, we first examined its expression in human myocardial tissue obtained at the time of LVAD implantation. Bulk RNA sequencing revealed significantly higher *LRRN4CL* expression in non-responders compared with responders and non-failing donor hearts. These findings were independently confirmed by RT-qPCR (**Figure 3D**), which demonstrated markedly elevated *LRRN4CL* transcript levels in the NR myocardium, while expression in R was comparable to non-failing donors.

To determine the cell types responsible for *LRRN4CL* expression, RNA in situ hybridization was performed on human cardiac tissue. The *LRRN4CL* signal was predominantly localized to cardiomyocytes and was markedly increased in NR myocardium compared with R and non-failing donors (**Figure 3E**). Collectively, these data reveal that increased *LRRN4CL* expression in cardiomyocytes is a feature of hearts less probable to experience LVAD-mediated myocardial recovery.

### LRRN4CL Localizes to the Sarcoplasmic Reticulum in Cardiomyocytes

LRRN4CL is predicted to be a single-pass transmembrane protein (31, UniProtKD, Q8ND94), but its subcellular localization in cardiomyocytes has not been previously defined. To determine the intracellular distribution of LRRN4CL, human iPSC-derived cardiomyocytes were transduced with an AAV9-aMHC-*LRRN4CL*-FLAG construct and examined by immunofluorescence microscopy (**Figure 3F**).

LRRN4CL-FLAG signal consistently demonstrated substantial co-localization with SERCA2, a marker of the sarcoplasmic reticulum (SR), indicating predominant localization within the SR compartment (**Figure 3F**). In contrast, minimal overlap was observed between LRRN4CL-FLAG and Na⁺/K⁺-ATPase, a marker of the sarcolemma, suggesting that cardiac LRRN4CL does not primarily localize to the plasma membrane. This intracellular distribution was not associated with overt changes in cellular morphology. These findings place LRRN4CL within the SR network of cardiomyocytes. Having established its subcellular localization, we next sought to define its functional role in cardiomyocytes.

### *LRRN4CL* Overexpression Induces Broad Transcriptional Remodeling in Human iPSC-Derived Cardiomyocytes

To determine how increased *LRRN4CL* expression alters cardiomyocytes, we again performed bulk RNA sequencing on human iPSC-CMs following adenoviral overexpression of *LRRN4CL* (**Figure 4A**). Cells were transduced with Ad-CMV-*LRRN4CL* at two increasing viral doses (MOI 20 and MOI 50) and compared with Ad-CMV-*GFP* controls.

**Figure 4.**
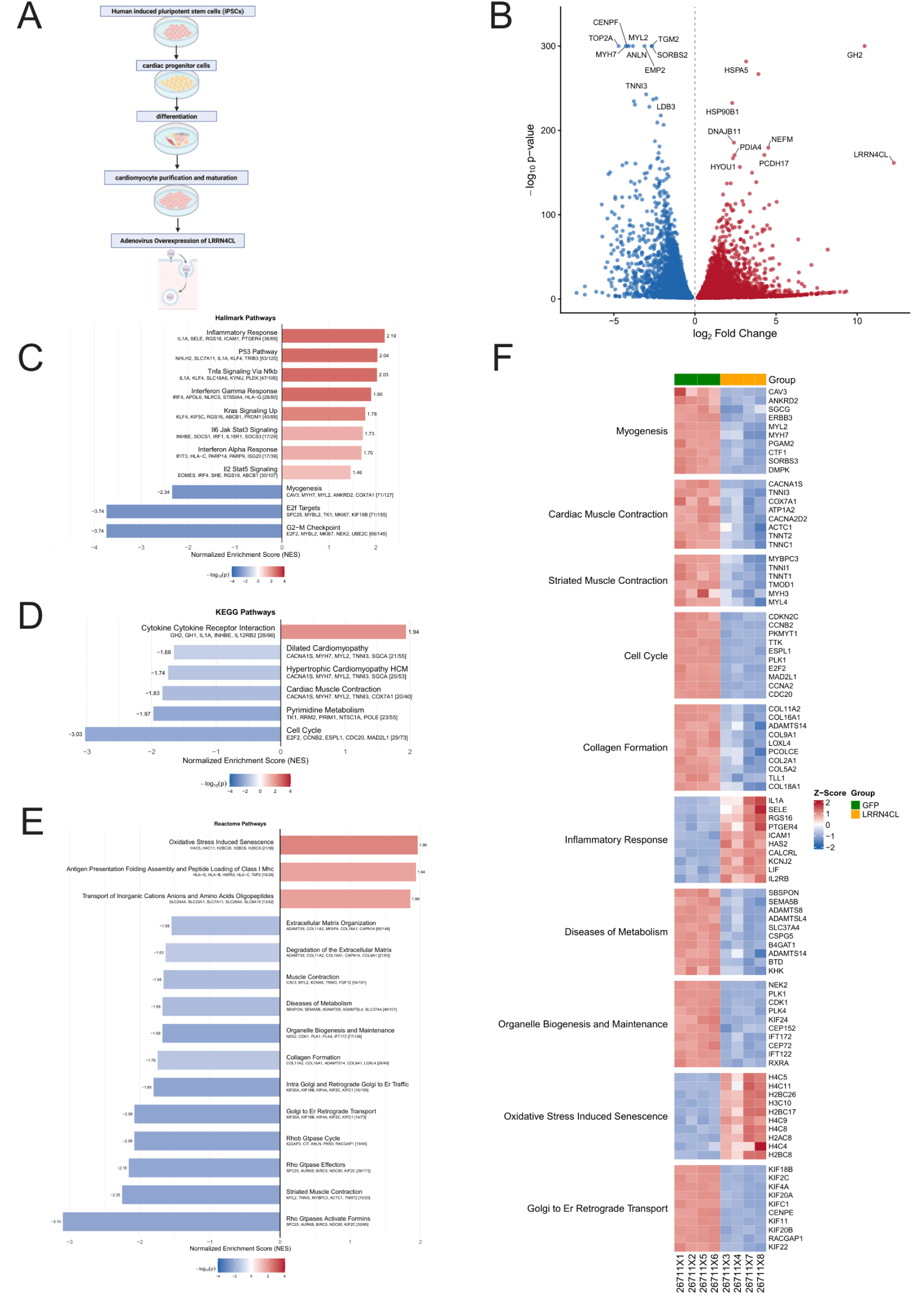
*LRRN4CL* modulates cardiac pathways that compromise structure and function. (**A**) Schematic overview of human iPSC–cardiomyocyte (iPSC-CM) differentiation and *LRRN4CL* overexpression timeline (n=4 for GFP and *LRRN4CL* overexpression groups). (**B**) Volcano plot showing DESeq2 differential expression results for LRRN4CL overexpression in iPSC-CMs (MOI 50). The plot represents the intersecting set of 6,771 genes significantly dysregulated at both MOI 20 and MOI 50, displayed using MOI 50 log₂ fold change and adjusted p-values. Significantly altered genes (padj<0.05) are highlighted, with selected genes labeled. Normalized enrichment score (NES) bar graph showing top enriched Hallmark (**C**), KEGG (**D**), and Reactome (**E**) gene sets from gene set enrichment analysis (GSEA) of differentially expressed genes. Bar colors reflect FDR-adjusted p-values (FDR<0.05). (**F**) Heatmap showing expression of genes from selected enriched pathways across GFP and *LRRN4CL* overexpression samples. Genes are grouped by pathways selected based on relevance GSEA significance (FDR<0.05). Colors indicate relative expression (row-scaled Z-scores).

Differential gene expression analysis revealed transcriptional changes following *LRRN4CL* overexpression. At the higher viral dose (MOI 50), 10,867 genes were differentially expressed relative to GFP controls, while 9,504 genes were differentially expressed at the lower dose (MOI 20). Intersection of these datasets identified 6,771 genes that were consistently dysregulated across both conditions, representing dose-independent transcriptional responses to *LRRN4CL* overexpression (**Figure 4B**). These shared differentially expressed genes were then used for pathway analysis, as they likely reflect direct biological consequences of increased *LRRN4CL* expression rather than secondary effects of viral load.

Gene set enrichment analysis (GSEA) of the shared differentially expressed genes highlighted pathways critical to cardiomyocyte structure and function, including significant downregulation of myogenesis (p=0.0019), cardiac muscle contraction (p=0.0224), and diseases of metabolism (p=0.0209) (**Figure 4C-F**). In parallel, *LRRN4CL* overexpression was associated with upregulation of stress associated pathways including inflammatory response (p=0.0018) and oxidative induced senescence (p=0.0054). Notably, enrichment of ER stress–associated pathways was concordant with our observed localization of LRRN4CL, suggesting a potential link between its intracellular distribution and downstream transcriptional stress responses. Together, these data demonstrate that increased *LRRN4CL* expression induces transcriptional remodeling in human iPSC-CMs, characterized by suppression of contractile and myogenic programs alongside activation of inflammatory, senescence, and stress pathways.

To further define regulatory programs driving the transcriptional remodeling induced by *LRRN4CL* overexpression, we performed transcription factor (TF) activity inference and gene regulatory network (GRN) analysis (**Supplemental Figure S11**). Using VIPER-based regulon analysis, multiple TFs demonstrated significant activity changes (FDR<0.05, |NES|>1.5), including regulators of metabolic adaptation (e.g., AHR, RARA, HNF4A), inflammatory and stress signaling which have canonically been associated with cardiovascular disease (e.g. TP53, FOS, NFKB1, E2F1), and contractile regulation (e.g. ZNF263). Integration of significantly regulated TFs and their top-ranked targets yielded a structured GRN highlighting central hubs such as NFKB1, TP53, E2F1, and MYC, linking inflammatory, metabolic, and proliferative pathways (**Supplemental Figure S11E**). Together, these data suggest that *LRRN4CL* overexpression drives coordinated shifts in regulatory networks governing stress adaptation in human iPSC-derived cardiomyocytes.

### Induction of Integrated Stress Response Transcription Factors Following *LRRN4CL* Overexpression

In human myocardial tissue, activating transcription factor 3 (*ATF3*) was identified as a transcript associated with adverse recovery phenotypes and was significantly upregulated in non-responders compared with responders and non-failing donor myocardium (**Supplemental Figure S4A**). ATF3 is a stress-inducible transcription factor commonly activated in response to cellular injury and maladaptive remodeling (32). Consistent with this observation, *LRRN4CL* overexpression in human iPSC-derived cardiomyocytes resulted in robust induction of *ATF3* (log2FC=1.51, FDR=6.97 × 10^⁻40^), along with significant upregulation of activating transcription factor 4 (*ATF4*; log2FC=0.86, FDR=1.28 × 10^⁻20^), a central mediator of the integrated stress response. ATF4 functions upstream of ATF3 within ER stress-associated signaling pathways, suggesting coordinated activation of stress-responsive transcriptional programs.

Given the localization of LRRN4CL to the SR compartment, the induction of *ATF4* and *ATF3* is consistent with activation of SR stress signaling. These findings provide mechanistic continuity between the stress-associated transcriptional signature observed in human myocardium and the cell-autonomous transcriptional response to *LRRN4CL* overexpression in cardiomyocytes, further supporting a model in which elevated *LRRN4CL* expression promotes SR stress and downstream maladaptive remodeling. This transcriptional signature provided a mechanistic basis for subsequent functional studies examining the impact of *LRRN4CL* overexpression on cardiomyocyte contractility and calcium handling.

### *LRRN4CL* Overexpression Impairs Contractile Dynamics in Human iPSC-CMs

To determine whether the transcriptional changes induced by *LRRN4CL* overexpression translate into functional impairment, we assessed contractile function in human iPSC-CMs. iPSC-CMs were imaged at baseline prior to viral transduction and again 72 hours following transduction with either Ad-CMV-*LRRN4CL* or Ad-CMV-*GFP* control virus.

*LRRN4CL* overexpression resulted in alterations in contractile kinetics compared with GFP controls (**Supplemental Video S1–4**). LRRN4CL-overexpressing cardiomyocytes showed reduced contraction amplitude (**Figure 5A–D**) and exhibited a prolonged contraction–relaxation cycle (**Figure 5E**). Quantitative analysis confirmed increased contraction–relaxation time (median difference 0.267s; 97.14% CI: 0.111 to 0.405s; p=0.0286), along with reduced contraction speed (−0.937µm/s; −1.307 to −0.671µm/s; p=0.0286) and reduced relaxation speed (−0.803µm/s; −1.017 to −0.479µm/s; p=0.0286) in LRRN4CL-overexpressing cells, indicating impaired contraction–relaxation kinetics (**Figure 5E–G**). These effects were consistent across replicates (n=4 per group), with 42 waveforms analyzed for LRRN4CL and 103 for GFP.

**Figure 5.**
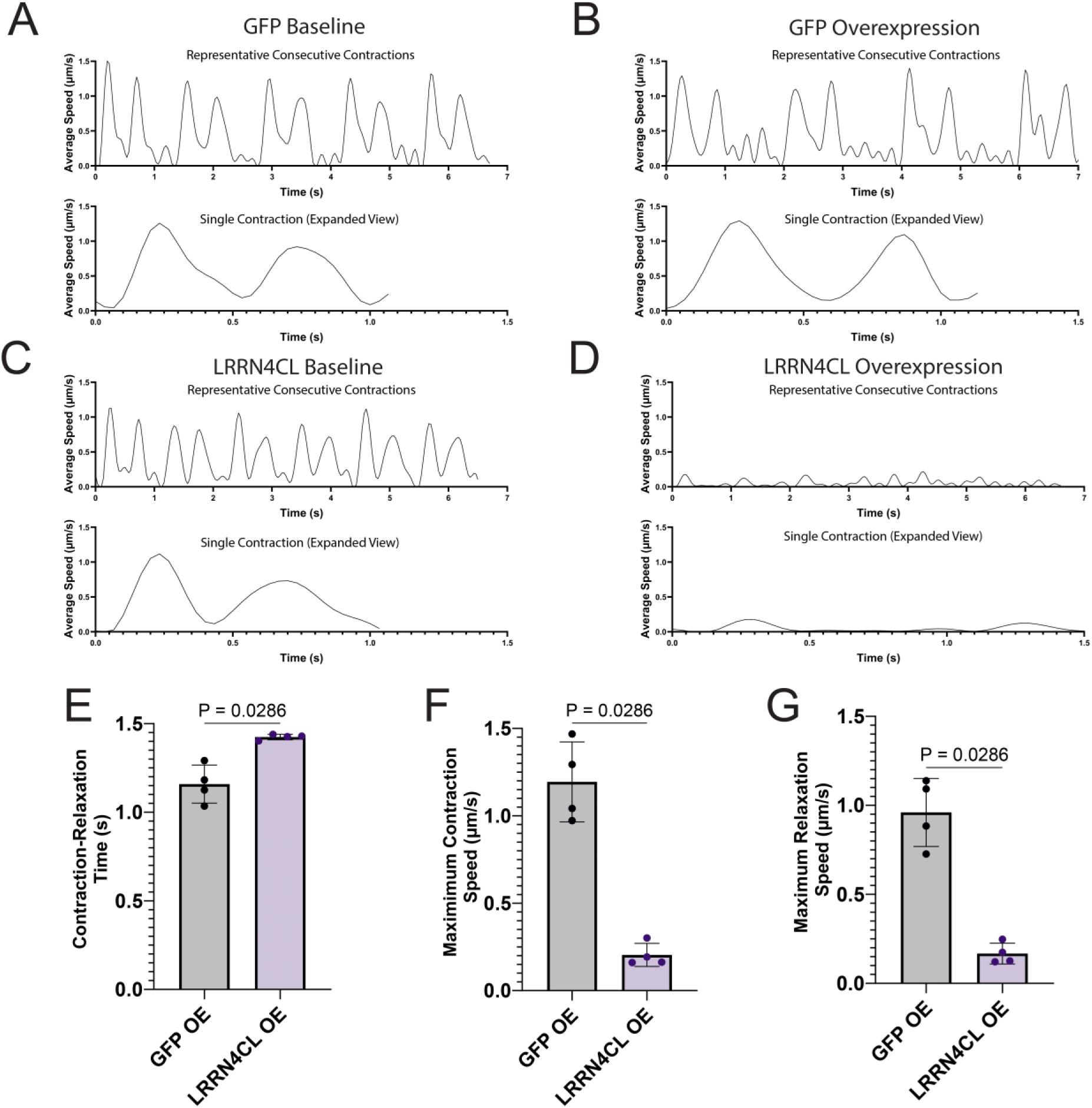
*LRRN4CL* overexpression impairs contractile dynamics in iPSC-CMs. **(A-D)** The top panels display consecutive representative contraction–relaxation waveforms. The bottom panels show a magnified view of a single representative waveform. Each waveform reflects the averaged contraction speed (µm/s) of all iPSC-CMs within the field of view during a single contraction–relaxation cycle for (**A**) GFP baseline, (**B**) GFP OE, (**C**) *LRRN4CL* baseline, and (**D**) *LRRN4CL* OE conditions. Traces illustrate changes in contraction amplitude and kinetics following *LRRN4CL* overexpression. Box plots show quantification of contractile parameters comparing GFP OE and *LRRN4CL* OE samples including (**E**) contraction–relaxation time, (**F**) maximum contraction speed, and (**G**) maximum relaxation speed. Contractile properties were quantified using CONTRACTIONWAVE (n=4 replicates per group), and waveforms were averaged per replicate. A total of 42 waveforms were analyzed for LRR groups and 103 for GFP groups. Statistical comparisons were performed using two-tailed Mann–Whitney tests. OE, overexpression.

### *LRRN4CL* Overexpression Impairs Cytosolic and Mitochondrial Calcium Handling in Human iPSC-Derived Cardiomyocytes

Given the impairment in contraction-relaxation kinetics and the localization observed following *LRRN4CL* overexpression, we next investigated whether altered calcium handling underlies the observed dysfunction. Cytosolic and mitochondrial calcium dynamics were assessed in human iPSC-CMs using live-cell confocal imaging.

Seventy-two hours following adenoviral transduction, iPSC-CMs were imaged under baseline conditions and following caffeine stimulation (20mM) to induce rapid SR calcium release. In *LRRN4CL*-overexpressing iPSC-CMs, caffeine-induced cytosolic calcium transients exhibited significantly prolonged decay kinetics compared with untreated (UT) and *LacZ*-expressing controls, with increased decay time relative to UT (median difference 1.6s; p=0.0127) and LacZ OE (6.8s; p<0.0001) (**Figure 6A, D**). These findings indicate impaired calcium reuptake or clearance following SR release.

**Figure 6.**
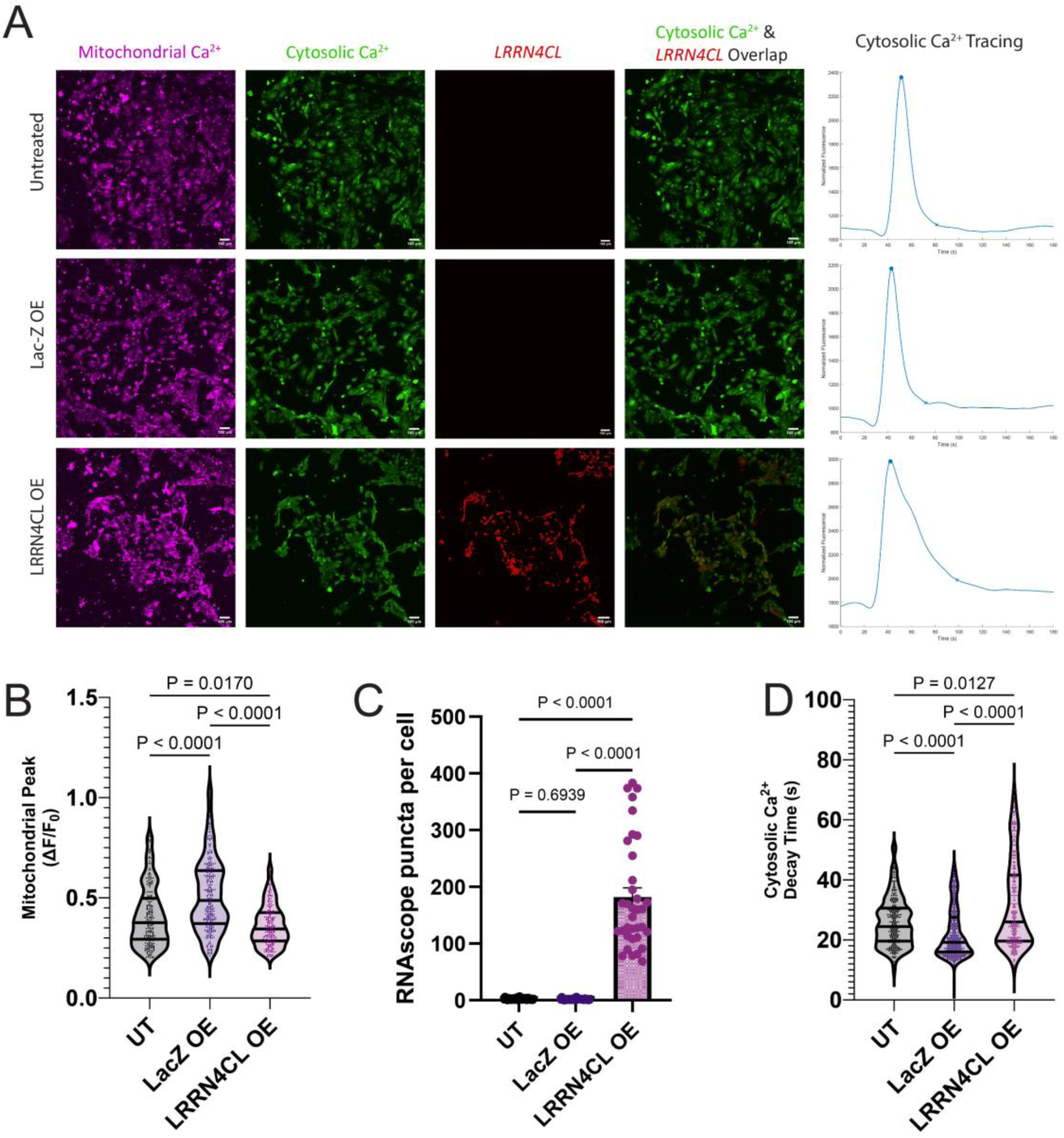
*LRRN4CL* overexpression alters cytosolic and mitochondrial Ca²⁺ dynamics in iPSC-CMs. (**A**) Representative images showing mitochondrial Ca²⁺ (purple), cytosolic Ca²⁺ (green), *LRRN4CL* RNA (red), and merged cytosolic Ca²⁺ and *LRRN4CL* RNA overlays in untreated (UT), *LacZ* overexpression (OE), and *LRRN4CL* OE conditions. Right panels show representative cytosolic Ca²⁺ traces post caffeine stimulation (20mM). (**B**) Quantification of mitochondrial Ca²⁺ peak amplitude (ΔF/F₀) (n=3 replicates per group; total cells analyzed: UT, 232; LacZ, 282; LRR, 195) (**C**) Quantification of *LRRN4CL* RNA puncta per cell (n=35 cells per group). (**D**) Quantification of cytosolic Ca²⁺ decay time (n=3 replicates per group; total cells analyzed: UT, 281; LacZ, 364; LRR, 267). Violin plots are presented as median with interquartile range. Median cytosolic Ca²⁺ decay times were 24.4s (IQR 19.6–30.6s) for UT, 19.2s (16.0–27.5s) for LacZ OE, and 26.0s (19.6–41.6s) for LRRN4CL OE. Median mitochondrial Ca²⁺ peak values were 0.376 ΔF/F₀ (IQR 0.293–0.498 ΔF/F₀) for UT, 0.487 ΔF/F₀ (0.372–0.636 ΔF/F₀) for LacZ OE, and 0.345 ΔF/F₀ (0.285–0.426 ΔF/F₀) for LRRN4CL OE. Statistical comparisons were performed using Kruskal–Wallis tests with Dunn’s multiple-comparisons correction.

In parallel, mitochondrial calcium handling was also altered by *LRRN4CL* overexpression. *LRRN4CL*-overexpressing iPSC-CMs exhibited significantly reduced mitochondrial calcium uptake following caffeine stimulation, with lower peak amplitude relative to UT (median difference −0.031ΔF/F₀; p=0.017) and LacZ OE (−0.142ΔF/F₀; p<0.0001) (**Figure 6B**). These findings suggest impaired coupling between cytosolic calcium transients and mitochondrial calcium buffering.

To confirm that calcium imaging was performed in cells expressing *LRRN4CL*, RNA in situ hybridization was conducted following live-cell imaging. *LRRN4CL* puncta were readily detected in imaged cells, and quantification confirmed robust expression in *LRRN4CL*-overexpressing cardiomyocytes relative to controls (**Figure 6A, C**).

Together, these data demonstrate that *LRRN4CL* overexpression disrupts calcium cycling, including delayed cytosolic calcium clearance and impaired mitochondrial calcium uptake. These abnormalities provide explanation for the prolonged contraction-relaxation kinetics and reduced mechanical efficiency observed in *LRRN4CL*-overexpressing cardiomyocytes and are consistent with the localization of *LRRN4CL* to the SR compartment.

### *LRRN4CL* Overexpression Limits Mitochondrial Bioenergetic Reserve in iPSC-CMs

To determine the impact of *LRRN4CL* overexpression on mitochondrial function in iPSC-derived cardiomyocytes, we performed a Seahorse XF Mito Stress Test (**Figure 7A**). Basal respiration was not significantly different between groups (**Figure 7B**). ATP-linked respiration was also unchanged (**Figure 7E**), indicating that oxidative phosphorylation adequately supports basal energetic demand in *LRRN4CL* OE cells.

**Figure 7.**
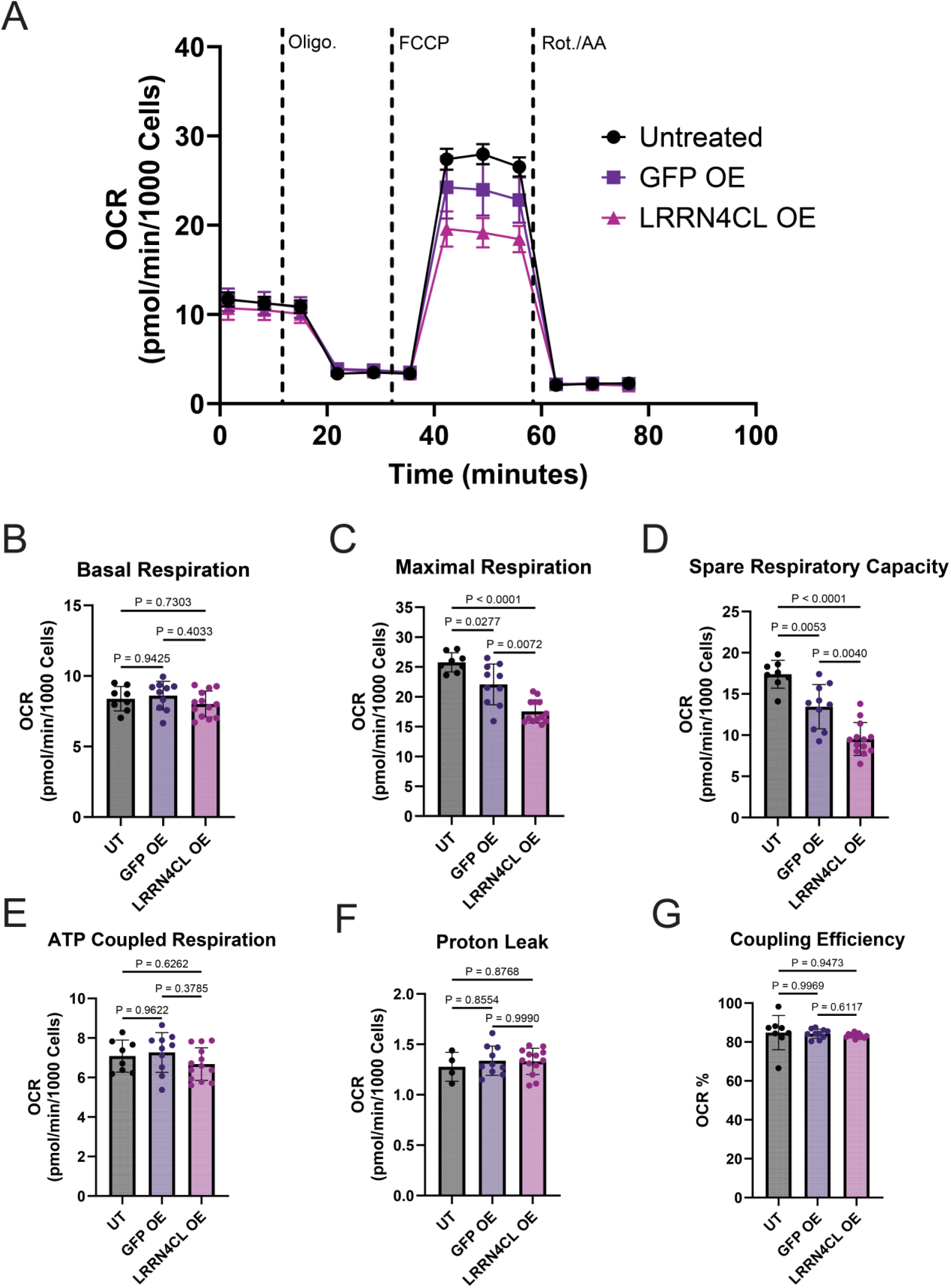
*LRRN4CL* overexpression impairs maximal mitochondrial respiration and spare respiratory capacity in iPSC-CMs. **(A)** Representative oxygen consumption rate (OCR) traces measured by Seahorse XF Cell Mito Stress Test in untreated (UT), GFP overexpression (GFP OE), and *LRRN4CL* overexpression (*LRRN4CL* OE) iPSC-CMs. Sequential injections of oligomycin (Oligo.), FCCP, and rotenone/antimycin A (Rot./AA) are indicated by dashed lines. Quantification of mitochondrial respiration parameters derived from the Mito Stress Test, including **(B)** basal respiration, **(C)** maximal respiration, **(D)** spare respiratory capacity, **(E)** ATP-linked respiration, **(F)** proton leak, and **(G)** coupling efficiency. *LRRN4CL* OE significantly reduced maximal respiration and spare respiratory capacity compared with UT and GFP OE controls, without affecting basal respiration, ATP-linked respiration, proton leak, or coupling efficiency. Data are presented as mean±SD (n=8 [UT], n=10 [GFP OE], and n=13 [LRRN4CL OE]). Some wells were excluded based on Seahorse quality control criteria. Statistical comparisons were performed using Welch’s ANOVA with Dunnett’s T3 multiple-comparisons test.

In contrast, maximal respiration was significantly reduced by LRRN4CL overexpression, with lower values compared to UT (mean difference −8.216 pmolO₂/min/1000cells; 95% CI −6.159 to−10.27; p<0.0001) and GFP OE (−4.514 pmolO₂/min/1000cells; −1.238 to −7.789; p=0.0072) (**Figure 7C**). This reduction resulted in a marked decrease in spare respiratory capacity, with LRRN4CL OE cells showing lower values compared to UT (−7.858 pmolO₂/min/1000cells;−5.708 to −10.01; p<0.0001) and GFP OE (−3.925 pmolO₂/min/1000cells; −1.238 to −6.612; p=0.0040) (**Figure 7D**).

Proton leak and coupling efficiency were unchanged (**Figure 7F, G**) across groups, indicating preserved mitochondrial membrane integrity and ATP production efficiency under basal conditions. Together, these findings demonstrate that *LRRN4CL* overexpression selectively impairs mitochondrial respiratory reserve capacity while maintaining basal mitochondrial function.

## DISCUSSION

Patients undergoing mechanical unloading and circulatory support with LVADs can experience structural and functional myocardial improvement to varying degrees, with a subset of patients improving to a degree that LVAD weaning can be considered (4–9,15). This clinical observation, combined with access to myocardial tissue from the LV apex at implantation, provides an unprecedented opportunity to characterize myocardial biology in advanced HF (15). In this multi-center study, we integrated pre-implant clinical characteristics with myocardial transcriptomics to identify predictors of myocardial recovery following LVAD-mediated unloading and circulatory support. To our knowledge, this is the first investigation to merge clinical and transcriptomic data using ML (21) to identify predictors of myocardial recovery in advanced HF. We found that established clinical features associated with reverse remodeling were recapitulated by our models, while *LRRN4CL* emerged as a molecular predictor of myocardial recovery assessed as a binary outcome (i.e. responder or non-responder) or assessed as continuous measures of functional and structural improvement.

Importantly, experimental validation using human cardiac tissue and iPSC-CMs demonstrated that increased *LRRN4CL* expression induces transcriptional remodeling, disrupts calcium handling, impairs contraction–relaxation kinetics, and produces bioenergetic inflexibility characterized by reduced mitochondrial reserve capacity. These findings provided a mechanistic link between ML-derived prediction and cardiomyocyte dysfunction suggesting that integrative clinical–molecular modeling can identify biologically meaningful predictors of recovery and may guide selection of patients who could benefit from a bridge-to-recovery strategy combining LVAD and standard HF pharmacological therapy.

Myocardial recovery following LVAD support is a heterogeneous clinical and biological phenomenon rather than a binary outcome (33,34). LVAD-mediated unloading leads to reduction in LV end-diastolic volume and pressure and circulatory support improves end-organ perfusion and reduces neurohormonal activation (34). These salutary effects lead to varying degrees of structural cardiac improvement with reductions in LV and left atrial volume and mass, and restoration of LV geometry (5,6,8,9,33,34,35). Although structural improvement and reductions of LV dimensions are almost universal, functional recovery varies, with LVAD-supported patients demonstrating various degrees of LVEF improvement (5,6,8,9). These findings highlight that myocardial recovery is a graded rather than a binary phenomenon and suggest potentially distinct biological processes driving structural and functional cardiac changes. Based on the above, we modeled recovery using both a binary clinical phenotype (R vs NR) and continuous measures of post-LVAD LVEF and LVEDD. Notably, *LRRN4CL* emerged as a consistent predictor across modeling strategies.

The ML framework identified clinical characteristics previously associated with LVAD-mediated recovery (6,8,9,16), including shorter HF duration, less LV chamber dilation, and higher pre-LVAD LVEF, consistent with prior observations suggesting that less burned-out cardiomyopathy might be associated with a greater reverse remodeling potential. While integrative models achieved the highest overall discrimination, performance was comparable to the parsimonious INTERMACS cardiac recovery (ICAR) clinical model (9), underscoring that the primary objective of the ML framework was biological feature discovery rather than incremental clinical predictive gain.

Integration of myocardial transcriptomics enabled identification of molecular features that may directly modulate cardiomyocyte adaptation to ventricular unloading and enhanced systemic circulatory support. *LRRN4CL* was enriched in cardiomyocytes from NR and localized predominantly to the SR compartment in iPSC-CMs, positioning it within intracellular networks central to calcium handling and excitation–contraction coupling. Although leucine-rich-repeat-containing proteins have been implicated in cardiac remodeling (36,37), the role of *LRRN4CL* in cardiomyocyte biology has not previously been defined.

Overexpression of *LRRN4CL* in human iPSC-CMs induced transcriptional remodeling characterized by suppression of myogenic and contractile programs and activation of SR stress and inflammatory pathways. These changes were accompanied by impaired cytosolic calcium clearance, reduced mitochondrial calcium uptake, and delayed contraction-relaxation kinetics, indicating disruption of calcium cycling. Seahorse analyses further demonstrated preserved basal and ATP-linked respiration but significantly reduced maximal respiration and spare respiratory capacity (SRC), indicating intact resting oxidative phosphorylation yet diminished ability to augment ATP production under stress. SRC is increasingly recognized as a critical determinant of cellular stress tolerance, reflecting the capacity of mitochondria to respond to acute energetic demand or injury (38,39). In cardiomyocytes, which operate at high energetic flux, even modest reductions in reserve capacity may limit contractile reserve before basal respiration declines (39).

Mechanistically, reduced mitochondrial reserve provides a unifying link between bioenergetic vulnerability and impaired calcium handling. Mitochondria and the SR are structurally coupled at contact sites that facilitate Ca²⁺ exchange and ATP delivery; mitochondrial ATP production fuels SERCA2a activity, and mitochondrial Ca²⁺ uptake modulates excitation–contraction efficiency (39,40). Thus, diminished SRC may become functionally limiting during increased workload or adrenergic stimulation, prolonging Ca²⁺ decay and impairing contraction–relaxation kinetics. Together, these findings support a *model* in which *LRRN4CL* overexpression induces bioenergetic inflexibility that compromises mitochondrial-SR coupling, contributing to impaired calcium handling and reduced contractile performance.

In addition to *LRRN4CL*, ML analyses identified distinct clinical and transcriptomic features associated with myocardial recovery defined as a binary outcome (R vs NR), as well as structural and functional cardiac improvement when modeled as a continuous metric. Across all models, clinical variables that have previously been associated with LVAD-mediated myocardial recovery were also identified in our study, including sex, HF symptoms duration, baseline LVEDD, LVAD configuration, and implementation of HF pharmacotherapy (6,8,9). Notably, clinical variables associated with structural versus functional improvement differed to a relatively large extent, reinforcing that structural and functional cardiac changes represent related but partially distinct mechanistic processes and clinical phenotypes. Additional transcripts (*RBMX2, EFCC1, TXNL4B, TCIM, KAZALD1*) were also identified (**Supplemental Figure S4**), implicating pathways related to RNA processing, calcium signaling, inflammation, and extracellular remodeling; expanded analyses are provided in the **Supplemental Results**. Future studies should investigate these additional predictors of recovery to clarify their mechanistic roles and potential translational and clinical relevance.

The identification and functional validation of *LRRN4CL* has translational implications for advanced HF management. The current models for clinical prediction of myocardial recovery in advanced HF incompletely capture the underlying myocardial biology. The specific findings of this study suggest that molecular information obtained before LVAD implantation may further refine risk stratification by identifying advanced HF patients whose myocardial biology is amenable to recovery following ventricular unloading and enhanced systemic circulatory support. In addition, the findings from this investigational platform of advanced HF could inform the prognosis and risk stratification of the broader population of patients with earlier stage HF.

From a translational point of view, this work illustrates how integrative clinical–transcriptomic approaches can move beyond clinical prediction toward identification of new and biologically precise therapeutic targets in HF following a bedside to bench and back approach.

Several limitations warrant consideration. Although this represents one of the largest myocardial transcriptomic datasets in LVAD-supported patients (n=208), the absolute number of patients achieving myocardial recovery is low, which limits feature stability and increases overfitting risk. Repeated cross-validation with a large training proportion was used to mitigate this risk, but future studies in independent cohorts will be important to further evaluate the generalizability of these findings. The two-stage feature selection strategy reduced dimensionality but may have favored transcripts with predominantly linear associations. Additionally, only pre-LVAD transcriptomics were analyzed, and dynamic molecular changes following unloading and circulatory support were not evaluated. Finally, functional validation in iPSC-CMs enables controlled mechanistic interrogation but does not fully recapitulate the maturity or complexity of adult myocardium.

In conclusion, the integration of human myocardial tissue studies, clinical phenotyping, machine learning-based prioritization, and functional validation establishes a biologically grounded framework for identifying predictors of myocardial recovery. In this study, *LRRN4CL* emerged as a novel molecular feature associated with lower likelihood of cardiac improvement across binary and continuous outcomes. Experimental validation demonstrated that increased *LRRN4CL* expression drives transcriptional remodeling, disrupts calcium handling, impairs contractility, and reduces mitochondrial reserve capacity, linking computational discovery to mechanistic insight. Collectively, these findings highlight the potential of integrative clinical–molecular modeling to (a) enrich the credibility of clinical outcome predictions with information reflecting underlying biology and (b) identify novel therapeutic targets for myocardial recovery applicable to broader HF patient populations.

## ACKNOWLEDGEMENTS

We thank Timothy Parnell for technical assistance and RNA Sequencing data analysis. We thank the Genomics core at the University of Utah for performing RNA sequencing and assisting with RT-qPCR. We thank the University of Utah Metabolic Phenotyping core for assisting with Seahorse metabolic assays. We acknowledge ChatGPT (OpenAI) was used to assist in drafting portions of this manuscript, and the authors take full responsibility for the accuracy, validity, and originality of the content. We also thank all the lab members and summer undergraduate trainees for sample collection and processing.

## SUPPLEMENTAL MATIERAL

Supplemental Methods

Supplemental Results

Tables S1–S4

Figures S1-S11

Videos S1–S4

References #41-53

## NON-STANDARD ABBREVIATIONS AND ACRONYMS

AAV9: Adeno-Associated Virus Serotype 9
AUC: Area Under the Curve
FDR: False Discovery Rate
FGSEA: Fast Gene Set Enrichment Analysis
GBM: Gradient Boosting Machine
GLM: Generalized Linear Model
GRN: Gene Regulatory Network
HF: Heart Failure
ICAR: INTERMACS Cardiac Recovery score
INTERMACS: Interagency Registry for Mechanically Assisted Circulatory Support
iPSC-CMs: Induced Pluripotent Stem Cell–Derived Cardiomyocytes
KEGG: Kyoto Encyclopedia of Genes and Genomes
LVAD: Left Ventricular Assist Device
LVEDD: Left Ventricular End-Diastolic Diameter
LVEF: Left Ventricular Ejection Fraction
MAE: Mean Absolute Error
ML: Machine Learning
MOI: Multiplicity of Infection
MSigDB: Molecular Signatures Database
NES: Normalized Enrichment Score
OCR: Oxygen Consumption Rate
OE: Overexpression
PRAUC: Precision-Recall Area Under the Curve
RF: Random Forest
ROC: Receiver Operating Characteristic
RT-qPCR: Reverse Transcription Quantitative Polymerase Chain Reaction
SERCA: Sarco/Endoplasmic Reticulum Calcium ATPase
SVM: Support Vector Machine
UT: Untreated

